# LINC complexes are mechanotransducers that discriminate Epithelial-Mesenchymal Transition programs

**DOI:** 10.1101/744276

**Authors:** Théophile Déjardin, Pietro Salvatore Carollo, Patricia M. Davidson, Cynthia Seiler, Damien Cuvelier, Bruno Cadot, Cecile Sykes, Edgar R. Gomes, Nicolas Borghi

## Abstract

LINC complexes are transmembrane protein assemblies that physically connect the nucleo- and cytoskeletons through the nuclear envelope. Dysfunctions of LINC complexes are associated with pathologies such as cancer and muscular disorders. The mechanical roles of LINC complexes in these contexts are poorly understood. To address this, we used genetically encoded FRET biosensors of molecular tension in LINC complex proteins of fibroblastic and epithelial cells in culture. We exposed cells to mechanical, genetic and pharmacological perturbations, mimicking a range of physiological and pathological situations. We show that LINC complex proteins experience tension generated by the cytoskeleton and act as mechanical sensors of cell packing. Moreover, the LINC complex discriminates between inductions of partial and complete epithelial-mesenchymal transitions (EMT). We identify the implicated mechanisms, which associate nesprin tension sensing with α-catenin capture at the nuclear envelope, thereby regulating β-catenin transcription. Our data thus implicate that LINC complexes are mechanotransducers that fine-tune β-catenin signaling in a manner dependent on the Epithelial-Mesenchymal Transition program.

## Introduction

The cell nucleus is not only the repository of the genome but also the largest organelle in most cells, whose shape response to mechanical cues was shown close to a century ago (Chambers and Fell 1931). The cytoskeleton mechanically couples the nucleus to cell adhesion complexes such that extracellular mechanical cues can affect the position and shape of the nucleus (Maniotis, Chen, and Ingber 1997). Such mechanical coupling is provided by outer nuclear transmembrane proteins, nesprins, whose KASH domain interacts with inner nuclear transmembrane SUN proteins in the perinuclear space (Lombardi et al. 2011). The cytoplasmic domain of nesprins can bind to the cytoskeleton and the nucleoplasmic domain of SUNs to the nucleoskeleton to form the so-called LINC complex: Linker of Nucleoskeleton and Cytoskeleton (Crisp et al. 2006).

Mutations in, or loss of LINC complex proteins impair nuclear envelope integrity (Crisp et al. 2006), nucleus anchoring (Starr and Han 2002; Grady et al. 2005), signaling to the nucleus (Neumann et al. 2010), chromosome positioning (Chikashige et al. 2006), DNA repair (Swartz, Rodriguez, and King 2014), genome transcription (Alam et al. 2016) and replication (Wang et al. 2018), which impacts cell polarity, migration, division, or differentiation in a variety of contexts. More recently, disruption of the LINC complex was shown to impair the induction by extracellular mechanical cues of chromatin stretching and transcription (Tajik et al. 2016), and nuclear translocation of transcription co-factors (Driscoll et al. 2015; Elosegui-Artola et al. 2017; Uzer et al. 2018). Genetically encoded biosensors of tension in nesprins now exist (Arsenovic et al. 2016), and direct force application on Nesprins was shown to elicit nucleus-autonomous signaling that targets nucleus stiffness (Guilluy et al. 2014). Yet, it is still unclear whether the consequences of LINC complex disruption as above result from an impairment of mechanotransduction within the LINC complex itself. Alternatively, they could result from a mere loss of mechanostructural integrity or even of a non-mechanical function. Thus, the mechanistic determinants of LINC complex cellular functions and their involvement in the many associated diseases mostly remain to be discovered (Janin et al. 2017).

Here, we focused on nesprin-2 giant (nesprin2G), a nesprin involved in nucleus positioning during cell polarization in migrating fibroblasts (Luxton et al. 2010; Borrego-Pinto et al. 2012). Nesprin2G forms a complex with and regulates the nuclear localization of β-catenin (Neumann et al. 2010), a major transcription co-factor in several morphogenetic processes. We had previously shown that, upon induction of partial or complete epithelial-mesenchymal transition (EMT) (Jolly et al. 2017; Aiello et al. 2018), epithelial cell packing regulates β-catenin signaling (Gayrard et al. 2018). We hypothesized that the LINC complex could participate in this mechanical regulation.

We combined molecular tension microcopy (Gayrard and Borghi 2016) with mechanical, genetic and pharmacological perturbations of fibroblastic and epithelial cells in culture. We found that the LINC complex is mechanosensitive to cell packing. Moreover, nesprin2G tension increases upon induction of partial, but not complete EMT, thereby defining two mechanisms of β-catenin nuclear translocation. Upon induction of complete EMT, relaxed nesprin2G recruits α-catenin at the nuclear envelope, which results in nuclear translocation of both catenins. Upon partial EMT however, tensed nesprin2G does not recruit α-catenin and only β-catenin translocates to the nucleus. Finally, we found that α-catenin sequesters β-catenin in the nucleus in a transcriptionally less active form.

## Results

### The LINC complex is under cytoskeleton-dependent tension balanced by cell adhesion

To assess the tension exerted on the LINC complex, we generated a short variant of nesprin2G known to rescue the nucleus positioning function of the full-length protein (Luxton et al. 2010) with a tension sensor module (TSMod, (Grashoff et al. 2010)), consisting of a pair of fluorescent proteins flanking a polypeptidic spring (Fig. 1A). The tension sensor is inserted between the transmembrane domain and the neighboring spectrin repeat (SR). As a tension-less control, we designed a short variant bearing mutations in the Calponin Homology domain (CH) that impair interaction with the cytoskeleton (CH mutant) (Luxton et al. 2010) (Fig. 1A). As expected, colonies of MDCK and NIH 3T3 cells stably expressing the construct able to bind the cytoskeleton (CB) exhibited a FRET index significantly lower than that of the CH mutant, supporting that the CB construct is under mechanical tension due to specific interaction with the cytoskeleton (Fig. 1B, C).

**Figure 1.**
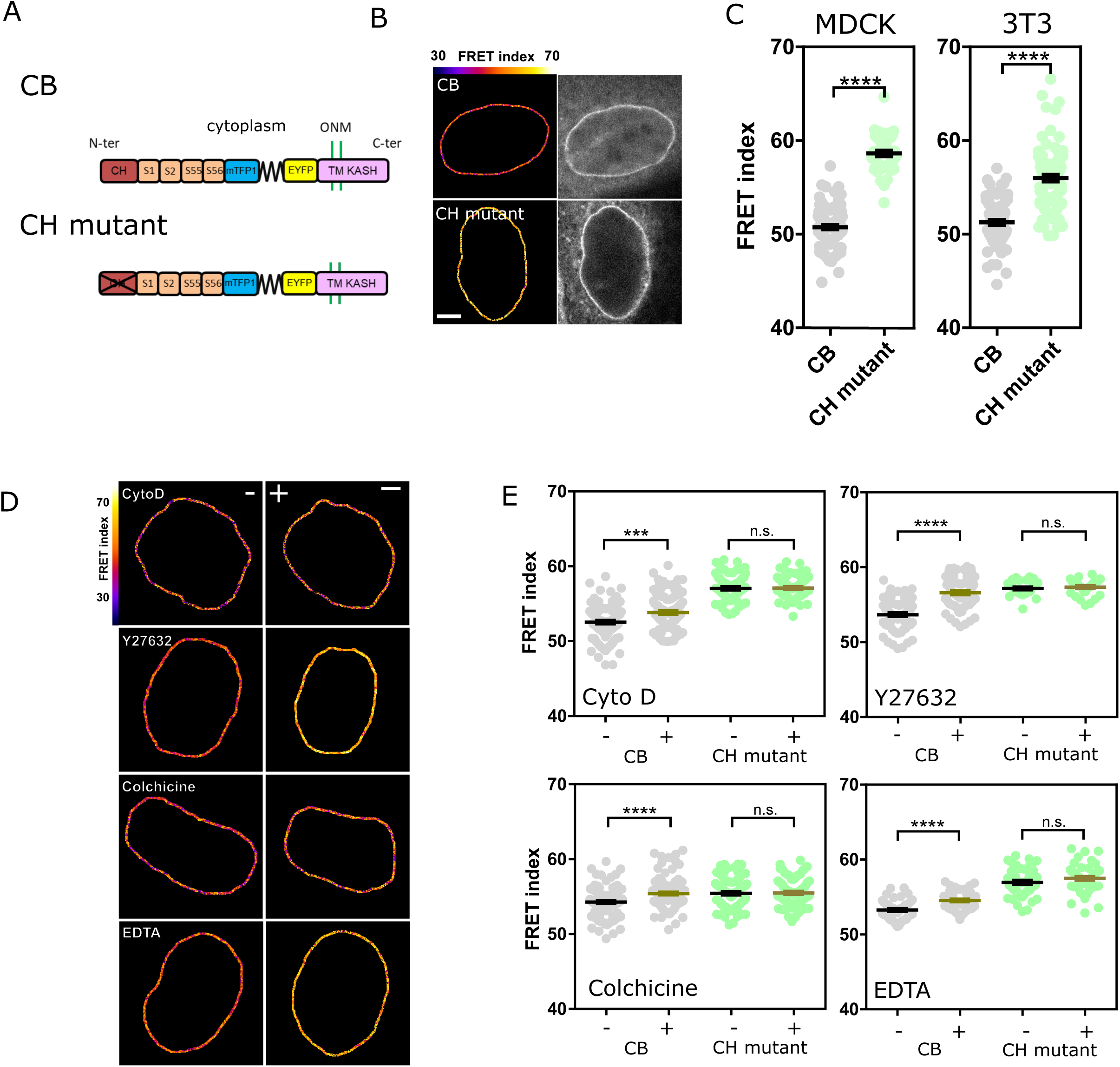
Nesprin 2 is under cytoskeleton-dependent tension and cytoskeleton-independent compression. A) Schematics of the Nesprin constructs. CB: cytoskeleton binding. CH: calponin homology domain. DNKASH: dominant negative KASH. ONM: Outer Nuclear Membrane. S1…S56: Spectrin repeat number. TM: Trans-Membrane. B) Typical nuclei expressing the Nesprin constructs above. Top: direct fluorescence. Bottom: FRET index map. C) FRET index of the Nesprin constructs above in MDCK (left) (n=97 CB, 45 CH mutant), and NIH 3T3 (right) (n=88 CB, 89 CH mutant) cells. D) FRET index map of the CB construct before and after pharmacological perturbations. Cyto D: cytoshalasin D. E) FRET index of the CB construct and the CH mutant before and 20 min after pharmacological perturbations) (n Cyto D = 108 CB, 71 CH mutant, n Y27632 = 74 CB, 30 CH mutant, n Colchicine = 112 CB, 88 CH mutant, n EDTA = 48 CB, 52 CH mutant). Bar=5 µm. Mean ± SEM. Krustal-Wallis (C) and Mann-Whitney (E) tests.

Moreover, the CB construct exhibited a significant FRET index increase in MDCK cells exposed to either the actin polymerization inhibitor cytochalasin D, the ROCK inhibitor Y27632 or the microtubule polymerization inhibitor colchicine (Fig. 1D, E). This supports that nesprin2G tension requires myosin contractile activity, and intact microtubule and filamentous actin networks. In contrast, the CH mutant did not exhibit significant FRET index changes in those conditions (Fig. 1D, E), indicating that disrupting actomyosin or microtubules does not further relieve tension when the cytoskeletal-binding domain is mutated. The tension generated by the cytoskeleton is thus exerted through the intact CH domain exclusively.

We next examined how the cytoskeleton-generated tension exerted onto nesprin2G was balanced. To do so, we perturbed calcium-dependent cell adhesion between MDCK cells by exposure to the calcium chelator EDTA. Impaired cell adhesion resulted in an increase of FRET index in the CB construct, but not in the CH mutant (Fig. 1D, E). This supports that cell adhesion is involved to balance the cytoskeletal tension on nesprin2G. We thus sought to assess nesprin sensitivity to extracellular mechanical cues.

### Tension on the LINC complex is sensitive to extracellular mechanical cues

To test nesprin2G sensitivity to extracellular mechanical cues, we first allowed MDCK and 3T3 cells to individually migrate through an array of obstacles with constrictions smaller than the nucleus diameter (Davidson et al. 2015). As cells migrated through constrictions, nucleus squeezing was accompanied by an increase in FRET index of the CB construct in nuclear regions within constrictions compared with regions outside constrictions (Fig. 2A, B). In contrast, the CH mutant exhibited an increase in FRET within constrictions that was not statistically significant. Therefore, these results support that cell migration through narrow constrictions reduces cytoskeleton-generated tension on nesprin2G in the region that is squeezed.

**Figure 2.**
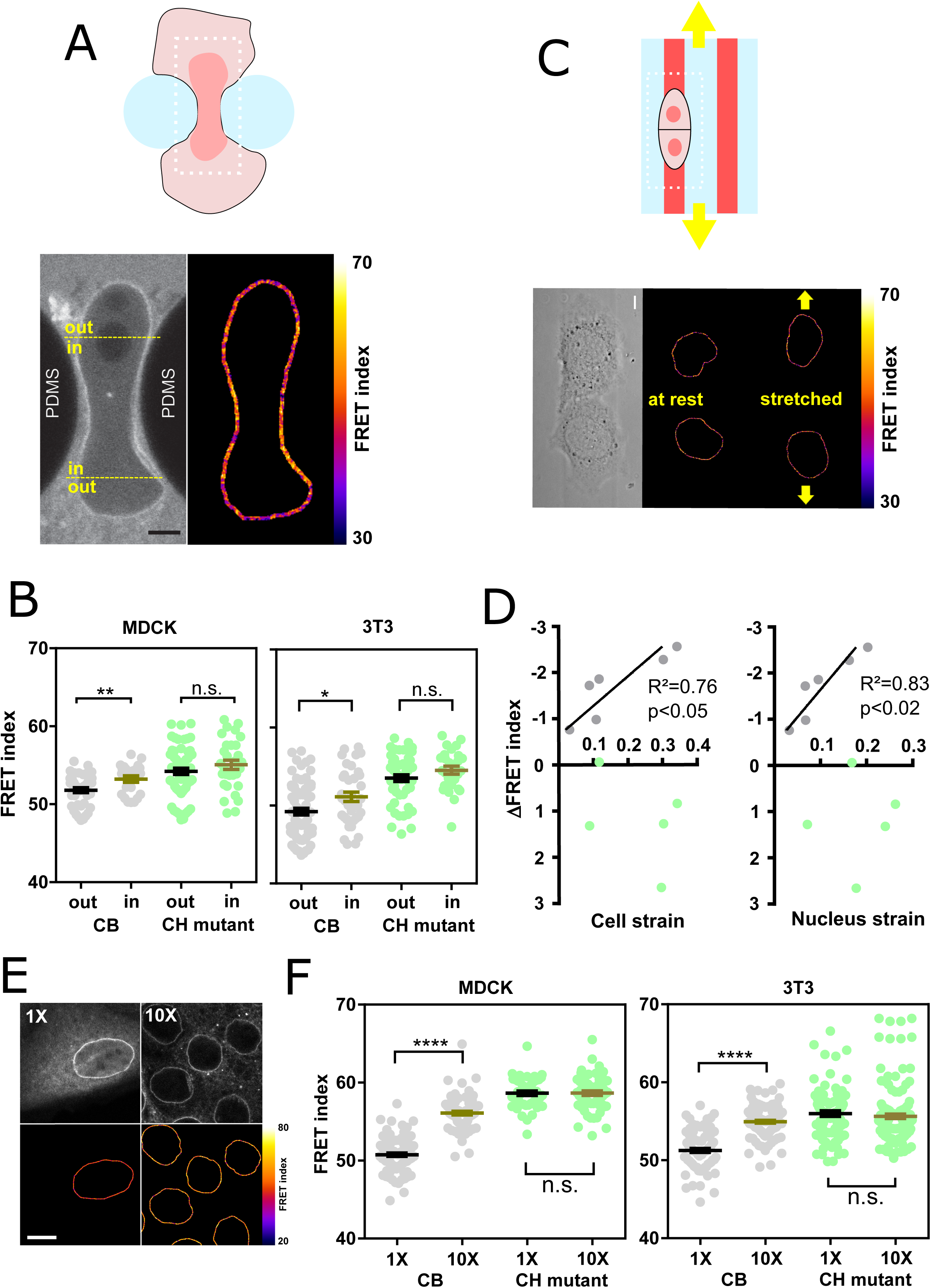
Nesprin 2 tension is sensitive to extracellular compression, stretch and cell packing. A) Top: Schematics of an event of cell migration through a narrow constriction. Bottom: direct fluorescence image and FRET index map from the dotted box above with boundaries between the region within the constriction and that outside. B) FRET index of the CB construct and the CH mutant inside and outside constrictions, in MDCK (left) (n= 24 CB, 40 CH mutant) and NIH 3T3 (right) (n= 48 CB out, 38 CH mutant) cells. C) Top: Schematics of the cell stretching experiment. Cells are plated on collagen stripes printed on a transparent, elastomeric sheet stretched in the direction of the adhesive stripes. Bottom: direct fluorescence image and FRET index map from the dotted box above. D) FRET index change upon stretching of the CB construct and the CH mutant as a function cell and nucleus strains. (n= 6 CB, 5 CH mutant). Solid lines are linear fits. E) MDCK cells expressing the CB construct plated at 5.10^2^ cells/mm^2^ (1X) and 5.10^3^cells/mm^2^ (10X). Top: fluorescence, bottom: FRET index map. F) FRET index of the CB construct and the CH mutant at 1X and 10X densities, in MDCK (left) (n= 97 CB 1X, 82 CB 10X, 45 CH mutant 1X, 61 CH mutant 10X) and NIH 3T3 (right) (n= 88 CB 1X, 120 CB 1X, 89 CH mutant 1X, 152 CH mutant 10X) cells. Bar=5 µm. Mean ± SEM. Mann-Whitney tests.

Next, we tested whether nesprin2G tension was sensitive to cell substrate stretching. To do so, we plated MDCK cells on adhesive patterns at the surface of a stretchable elastomer sheet substrate (Fink et al. 2011). Upon uniaxial substrate stretching, the FRET index in the CB construct significantly decreased in proportion to both cell and nucleus strains, while it did not in the CH mutant (Fig. 2C, D). Thus, nesprin2G tension is sensitive to cell substrate stretching through the cytoskeleton.

Altogether, these results show that nesprin2G tension is sensitive, through the cytoskeleton, to extracellular deformations both in compression and stretch. We thus hypothesized that nesprin2G could sense cell packing.

### Cells sense cell packing within the nucleus through the cytoskeleton and the LINC complex

To test how nesprin2G responds to cell packing, we compared cells plated at low density in colonies as in previous experiments (5.102 cells/mm2) with cells at confluence (5.10^3^ cells/mm2). In cells at confluence, the nucleus cross-sectional area was much smaller (Fig. S1A), and the CB construct exhibited a much higher FRET than in cells at low density, while the CH mutant did not (Fig. 2E, F). Tension on nesprin2G is thus dependent on cell density, in a manner that depends on cytoskeleton binding.

To assess whether this sensitivity propagated within the nucleus, we measured the FRET index in a TSMod tension sensor inserted between the transmembrane and nucleoplasmic domains of SUN2, such that it reported forces between the inner nuclear membrane and the nucleoskeleton (Fig. S1B). In this construct, the FRET index was higher overall than in the CB construct at equivalent cell density, suggesting a higher level of compression of the sensor not inconsistent with a highly crowded nucleoplasm. Moreover, cell confluence increased the FRET index in this construct compared to cells at low density, consistent with an increase in compression or a release of tension between the inner nuclear membrane and the nucleoskeleton (Fig. S1B). Thus, these results support that cell packing is sensed within the nucleus through the cytoskeleton and the LINC complex: the lower the packing, the higher the tension on both sides of the LINC complex.

### Tension in the LINC complex responds to cell packing upon induction of partial EMT

Epithelial sheet wounding results in decreased cell packing at the wound edge, where cells adopt a mesenchymal-like phenotype yet migrate as a cohesive group, a model of partial EMT (Gayrard et al. 2018). Thus, we hypothesized that tension in the LINC complex would increase at the edge of a wounded sheet. In agreement, the FRET index from the CB construct exhibited in wounded MDCK and 3T3 sheets a positive gradient from the wound edge back, indicative of a tension at the wound edge higher than in the back of the monolayer (Fig. 3A, B). Edge cells did not exhibit a significant difference in FRET index between the front and back of their nuclei (Fig. S1C). Of note, the CH mutant also exhibited some FRET index decrease toward the wound edge in MDCK (but not NIH 3T3), although to a smaller extent than the CB construct, suggesting a release of some compression independent of cytoskeleton binding in this condition.

**Figure 3.**
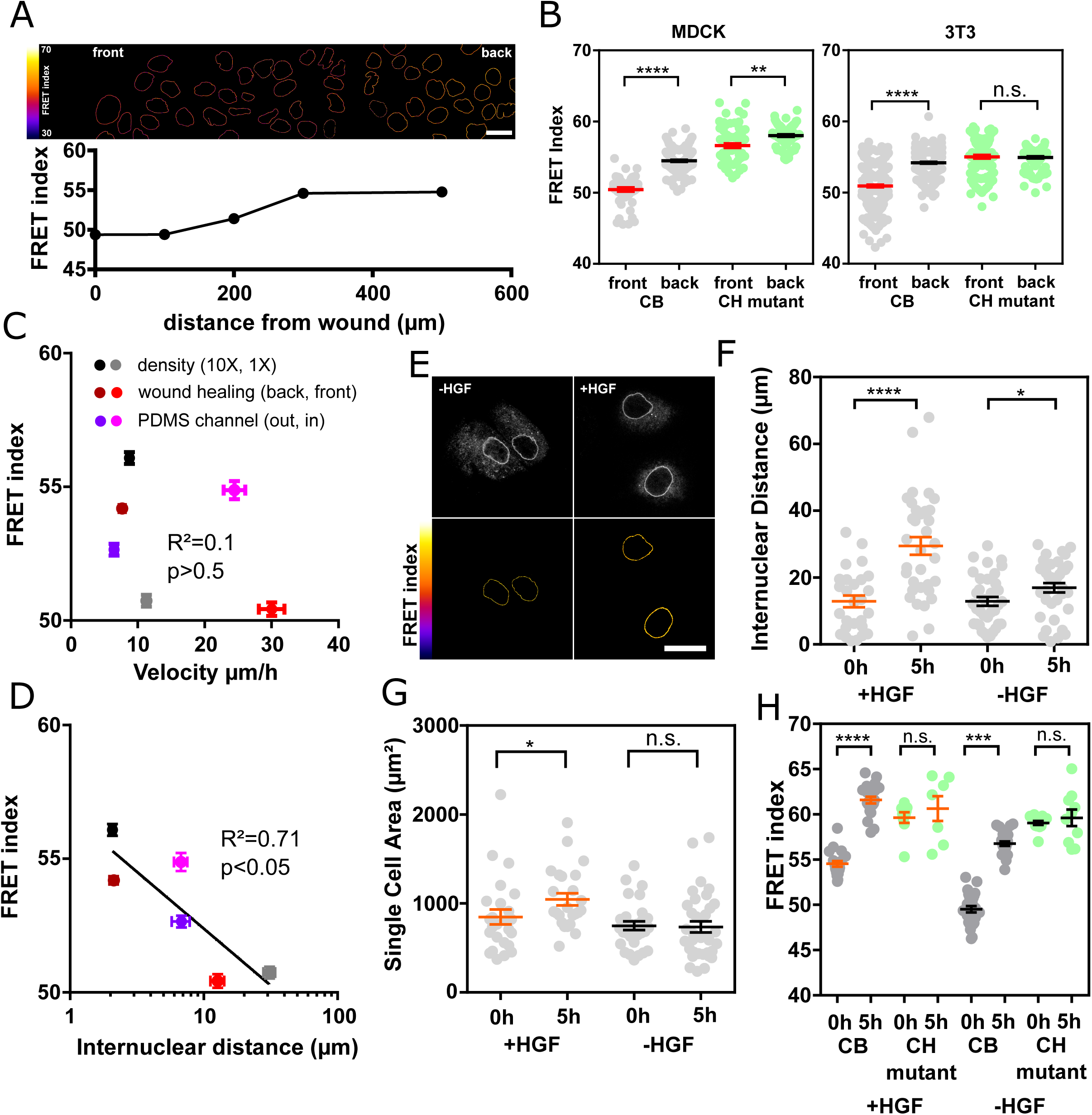
Nesprin 2 tension is differentially sensitive to induction of partial and complete EMT. A) Top: FRET index map of a wounded MDCK monolayer expressing the CB construct. Bottom: FRET index as a function of the distance from the front. B) FRET index of the CB construct and the CH mutant at the front and back (500 µm) of the monolayer, in MDCK (left) (n= 70 CB front, 130 CB back, 73 CH mutant front, 91 CH mutant back) and NIH 3T3 (right) (n= 363 CB front, 311 CB back, 125 CH mutant front, 143 CH mutant back) cells. Solid line to guide the eye. C) FRET index of the CB construct in MDCK cells as a function of cell migration velocity from experiments at low and high cell densities, upon epithelial wounding, and collectively migrating from outside (out) to inside (in) 40µm-wide channels) (n= 240 1X, 255 10X, 111 front, 110 back, 64 in, 78 out). D) FRET index of the CB construct in MDCK cells as a function of internuclear distance from experiments at low and high cell densities, upon epithelial wounding and upon entering 40µm-wide channels. Solid line is the linear fit (n= 22 1X, 22 10X, 22 front, 22 back, 33 in, 18 out). E) Direct fluorescence and FRET index maps of the CB construct in MDCK cells after 5hrs with or without HGF. F) Internuclear distance of CB construct-expressing MDCK cells through time with or without HGF addition (n +HGF= 26 0h, 27 5h, n –HGF= 30 0h, 32 5h). G) Single cell area of CB construct-expressing MDCK cells through time with or without HGF addition (n +HGF= 25 0h, 24 5h, n –HGF= 29 0h, 35 5h). H) FRET index of the CB construct and CH mutant in MDCK cells through time with or without HGF addition (n +HGF= 20 CB 0h, 22 CB 5h, 9 CH mutant 0h, 7 CH mutant 5h, n −HGF= 26 CB 0h, 26 CB, 5h, 13 CH mutant 0h, 10 CH mutant 5h). Bar=20 µm. Mean ± SEM. Mann-Whitney tests.

To assess whether increased cytoskeletal tension on nesprin2G is due to reduction in cell packing and not to increased cell migration velocity, we compared inter-nuclear distance and cell velocity with FRET in cells at the edge and the bulk of a wounded monolayer, in cells at low density and confluence as above, and in cells collectively migrating within 40 µm-wide channels that maintain cell density while allowing cell migration (Fig. 3C, D). From all these conditions, we found that FRET index correlated with inter-nuclear distance but not with cell velocity (Fig. 3C, D).

Altogether these results support that induction of partial EMT in a wound healing model leads to increased tension on the LINC complex due to decreased cell packing rather than cell migration.

### Tension in the LINC complex does not respond to cell packing upon induction of complete EMT

Exposure of cell colonies to HGF, a model of complete EMT where cells eventually migrate individually, induces decreased cell packing in the first hours before cell dissociation (Gayrard et al. 2018). Thus, we expected HGF exposure to induce an increase in tension in the LINC complex. To test this, we monitored how the inter-nuclear distance, the cell spread area, and the FRET index of nesprin2G constructs in MDCK colonies changed over time with or without exposure to HGF. Cells exposed to HGF exhibited a significantly larger increase in inter-nuclear distance and cell spread area than non-exposed cells over the same time frame (Fig. 3E-G), which confirms that HGF decreases cell packing. Unexpectedly, the FRET index of the CB, but not the CH construct exhibited an increase over time, whether colonies where exposed to HGF or not (Fig. 3E, H). These results reveal that the cytoskeletal tension exerted on the LINC complex has not reached a steady-state in cell colonies and slowly relaxes over time. Additionally, the lack of difference with or without HGF shows that tension in the LINC complex is not responsive to cell packing upon induction of complete EMT.

Thus, the LINC complex exhibits distinct mechanical responses to cell packing whether it results from induction of partial or complete EMT. While the causes for this differential response may be the focus of future studies, here we focused on its link with EMT-related signaling.

### Nesprin cytoplasmic domain defines two mechanisms of β-catenin nuclear translocation differentially activated upon induction of partial or complete EMT

We hypothesized that the distinct mechanical responses of nesprin2G upon induction of partial and complete EMT associated with a differential regulation of β-catenin signaling. To test this, we assessed changes in the nucleo/cytoplasmic balance of α-catenin-GFP and β-catenin-GFP upon HGF stimulation of MDCK cells. HGF stimulation induced β-catenin nuclear translocation, as expected (Gayrard et al. 2018), but also that of α-catenin (Fig. 4A, B). This is in stark contrast with the behavior of α-catenin in cells induced to undergo partial EMT in wound healing experiments, where the release of catenins from the plasma membrane in leader cells results in nuclear translocation of β-catenin but the retention of α-catenin in the cytoplasm (Gayrard et al. 2018). Moreover, we showed that β- and α-catenin nuclear contents were reduced to levels indistinguishable from that of unstimulated cells (Fig. 4A, B) in cells transiently expressing mCherry-DNKASH, a truncated nesprin that lacks the cytoplasmic domain and displaces endogenous nesprins from the nuclear envelope (Fig. S2A) (Luxton et al. 2010). To assess the consequences on β-catenin-dependent transcription, we transiently expressed mCherry-DNKASH in a TOPdGFP cell line, in which GFP expression is under the control of β-catenin transcriptional activity (Maher et al. 2009). These cells were exposed to the β-catenin degradation inhibitior LiCl for 10hrs to increase β-catenin levels. In control cells LiCl exposure resulted in GFP expression (Fig. S2B), consistent with excess β-catenin accumulation in the nucleus (Gayrard et al. 2018). In mCherry-DNKASH expressing cells, decreased GFP levels indicated decreased β-catenin transcriptional activity (Fig. S2B), consistent with the loss of nuclear β-catenin upon the same perturbation (Fig. 4A).

**Figure 4.**
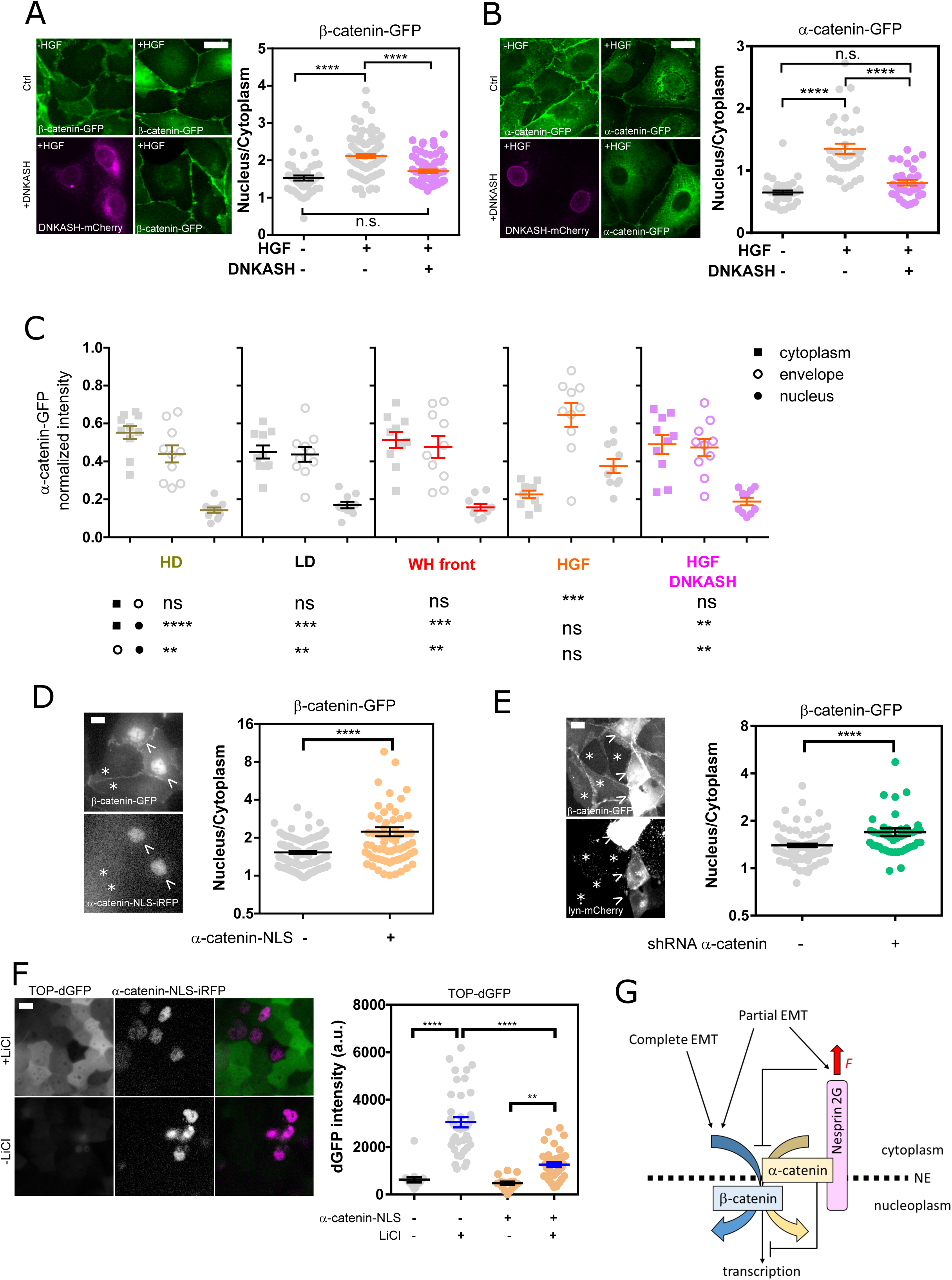
Nesprin 2 regulates catenins nuclear translocation. A) Left: MDCK cells stably expressing b-catenin-GFP with and without mCherry-DNKASH, with and without HGF addition. Right: b-catenin nucleus/cytoplasmic balance (GFP intensity ratio) as a function of HGF and DNKASH (n= 43--, 89+-, 65++). B) Left: MDCK cells stably expressing a-catenin-GFP with and without mCherry-DNKASH, with and without HGF addition. Right: a-catenin nucleus/cytoplasmic balance (GFP intensity ratio) as a function of HGF and DNKASH (n= 35 --, 36 +-, 31 ++). C) Relative cytoplasmic, nuclear envelope, and nuclear levels of a-catenin in cells plated at high density (HD, 10X), low density (LD, 1X), at the wound front, upon HGF exposure, and upon HGF exposure and mCherry-DNKASH expression. n=10 cells for each condition and compartment. D) Left: MDCK cells stably expressing b-catenin-GFP with transient expression of NLS-iRFP-a-catenin (arrow) and without (star). Right: b-catenin nucleus/cytoplasmic balance (GFP intensity ratio) as a function of NLS-iRFP-a-catenin expression (n= 112 -, 66 +). E) Left: MDCK cells stably expressing b-catenin-GFP with transient expression of shRNA against a-catenin and lyn-mCherry (arrow) and without (satr). Right: b-catenin nucleus/cytoplasmic balance (GFP intensity ratio) as a function of shRNA/lyn-mCherry co-transfection (n= 102 -, 44 +). F) Left: MDCK cells stably expressing TOP-dGFP and transiently expressing NLS-iRFP-a-catenin after 10hrs with and without LiCl. Right: dGFP intensity as a function of NLS-a-catenin-iRFP expression and LiCl (n= 17 ---, 39 -+, 17 +-, 39 ++). G) Working model. See text for details. Bar=10 µm. Mean ± SEM. Kruskal-Wallis tests.

Altogether these results support that β-catenin nuclear translocation and subsequent transcriptional activity can occur through distinct mechanisms: in cells induced to undergo complete EMT, nesprin2G is relaxed and its cytoplasmic domain required for β- and α-catenin nuclear translocation and β-catenin activity, while in cells induced to undergo partial EMT in wound healing experiments, nesprin2G is tensed and β-catenin translocates into the nucleus alone.

### Relaxed, but not tensed nesprin2G recruits α-catenin to the nuclear envelope

We then assessed whether the cytoplasmic domain of nesprin2G, which contains a binding site for α-catenin in a complex with β-catenin (Neumann et al. 2010), contributed to the nuclear translocation of catenins by promoting the recruitment of α-catenin to the nuclear envelope. Consistent with this hypothesis, we found that in HGF-stimulated cells, α-catenin accumulated at the nuclear envelope even more than within the nucleus, and that both these accumulations were abolished in cells expressing mCherry-DNKASH, which lacks the binding site for α-catenin (Fig. 4B,C Fig. S2C). This suggests that α-catenin nuclear translocation requires its recruitment to the nuclear envelope by nesprins. Consequently, we hypothesized that increased tension on nesprin2G in cells induced to undergo partial EMT could explain the lack of α-catenin nuclear translocation by preventing its recruitment to the nuclear envelope. Consistently, we found no recruitment of α-catenin to the envelope compared to the cytoplasm in cells a the wound front (Fig. 4C, Fig. S2C).

Altogether, these results show that in cells induced to undergo complete EMT, relaxed nesprin2G recruits α-catenin to the nuclear envelope, while in cells induced to undergo partial EMT, tensed nesprin2G does not.

### Nuclear localization of α-catenin causes β-catenin nuclear retention, but in a transcriptionally less active form

To assess the role of nuclear α-catenin in β-catenin signaling, we transiently expressed NLS-iRFP-α-catenin in cells stably expressing β-catenin-GFP. Compared to control cells, NLS-iRFP-α-catenin cells exhibited increased nuclear β-catenin (Fig. 4D). Thus, nuclear α-catenin promotes β-catenin nuclear localization. To determine whether α-catenin is involved in β-catenin translocation or nuclear retention, we examined β-catenin localization in cells transiently expressing a shRNA against α-catenin (Capaldo and Macara 2007). These cells displayed higher nuclear β-catenin levels compared to controls (Fig. 4E). This implies that not only α-catenin is dispensable for β-catenin nuclear translocation, but also suggests that its extranuclear pool opposes constitutive β-catenin nuclear localization. Finally, we assessed the effects of nuclear α-catenin on β-catenin transcriptional activity. To do so, we transiently expressed NLS-iRFP-α-catenin in TOPdGFP cells exposed, or not, to LiCl for 10hrs to increase β-catenin levels. Cells not exposed to LiCl exhibited low levels of GFP regardless of NLS-iRFP-α-catenin expression, as expected for cells with basal β-catenin levels. In contrast, cells exposed to LiCl exhibited significantly higher GFP levels, as expected for cells with high β-catenin levels, but to a much lower extent in cells expressing NLS-iRFP-α-catenin (Fig. 4F). Thus, nuclear α-catenin limits β-catenin transcriptional activity. Altogether, these results support that α-catenin nuclear translocation favors β-catenin nuclear localization, but in a transcriptionally less active form.

## Discussion

In this work, we sought to determine whether and how the LINC complex participated in the mechanical regulation of β-catenin signaling during EMT. We found that nesprin2G tension increases during partial, but not complete EMT. Upon induction of complete EMT, relaxed nesprin2G recruits α-catenin at the nuclear envelope, which results in nuclear translocation of both catenins. Upon partial EMT however, tensed nesprin2G does not recruit α-catenin and β-catenin nuclear translocation occurs independently. Once in the nucleus, α-catenin sequesters β-catenin in a transcriptionally less active form.

Using instrument-specific FRET index to FRET efficiency calibration (Fig. S3A) and previously published FRET efficiency to force calibration (see Material & Methods), we estimate that forces exerted by the cytoskeleton on nesprin2G can be as high as 8pN, the full range of the force sensor (Fig. 1). Remarkably, perturbation of both the actomyosin machinery and the microtubule network affect nesprin2G tension. This is consistent with the actin- and microtubule-binding properties of CH-domains (Hayashi and Ikura 2003; Goldsmith et al. 1997). Conditions aimed at mimicking cell morphological changes in a range of physiological and pathological situations all result in cytoskeleton-dependent tension changes consistent with that previously observed in cell adhesion proteins (Grashoff et al. 2010; Borghi et al. 2012). The sensor response in SUN2 is consistent with a pre-compressed state (Fig. S1B). Using the same sensor, protein compression has previously been evidenced in vinculin at Focal Adhesions (Rothenberg et al. 2015; Sarangi et al. 2017) and in the glycocalyx protein MUC1 (Paszek et al. 2014). Our results are consistent with the sensor being sensitive to such a compression and suggest this occurs constitutively within the nucleus.

A substantial part of cytoskeleton-dependent nesprin tension is balanced cell-autonomously, as can be seen from individual cells migrating through narrow constrictions (Fig. 2). Focal adhesions, which anchor actin stress fibers to the extracellular matrix, are well positioned to play a role in this balance. Indeed, a number of adherent cells display a nesprin-dependent perinuclear actin cap with fibers terminated by FAs (Khatau et al. 2009; Chambliss et al. 2013; Kim et al. 2012), cell stretching with integrin ligand-coated beads results in cytoskeleton- and nesprin-dependent nucleus stretching (Maniotis, Chen, and Ingber 1997; Lombardi et al. 2011), and nucleus anchoring to the cytoskeleton affects cell-substrate traction forces (Shiu et al. 2018). Here, we bring a direct demonstration that nesprin2G tension responds to substrate mechanics: provided that nesprins can bind to the cytoskeleton through their CH domains, their tension increases in remarkable proportion with cell and nucleus strain upon stretching of the cell substrate (Fig. 2). While previous studies have mostly focused on isolated cells, we also show here how nesprin tension changes in a cell assembly. Remarkably however, this tension does not necessarily correlate with that in E-cadherins, as nesprin and E-cadherin tension gradients are opposite in cells undergoing partial EMT (Fig. 3 and (Gayrard et al. 2018)). This points to a force balance regulation that likely depends on all adhesion complexes and variable fractions of mechanically engaged proteins.

We show that cell packing is a critical determinant of tension in the LINC complex, on both sides of the nuclear envelope (Fig. 2, S1B). We thus bring a direct demonstration that the LINC complex is a bona fide mechanosensor of cell packing at the nuclear envelope. Nevertheless, a decrease in cell packing results in an increase in nesprin tension at the front of an epithelial monolayer in partial EMT, but does not upon induction of complete EMT by HGF (Fig. 3). This differential response remains unexplained, but it supports that the LINC complex is a mechanosensor able to discriminate between inductions of various EMT programs. This makes nesprin2G tension a better predictor of EMT program than E-cadherin tension or β-catenin nuclear localization. Whether this can be further harnessed in the context of diseases may be the focus of future investigations.

Over the last decade, a number of signaling pathways regulating proliferation have been found to respond to cell confinement, and to depend on the LINC complex, supporting a role of the latter in the former. Confinement modulates YAP/TAZ nuclear translocation and ERK activity cell-autonomously (Dupont et al. 2011; Logue et al. 2015) and in a multicellular context (Aragona et al. 2013; Aoki et al. 2013). Moreover, mechanical induction of YAP nuclear translocation requires the cytoplasmic domain of nesprins at the nuclear envelope (Elosegui-Artola et al. 2017). However, it does not always appear to require a contractile cytoskeleton (Driscoll et al. 2015), which questions whether nesprins bear an actual mechanical function in this process. In addition, cell packing modulates β-catenin signaling downstream of FAK (Gayrard et al. 2018), and mechanical induction of β-catenin nuclear translocation is impaired upon disruption of the LINC complex (Uzer et al. 2018). While nesprin2G can interact with catenins (Neumann et al. 2010), it was however unknown whether this interaction could be mechanically regulated. Here, we took advantage of the differential response of nesprin2G to partial and complete EMT to show that the relaxed cytoplasmic domain of nesprin2G is required for α-catenin recruitment at the nuclear envelope and catenin nuclear translocation, while tensed nesprin2G does not recruit α-catenin at the envelope nor does it allow its nuclear translocation (Fig. 3, 4). Thus, nesprin2G tension predicts nuclear translocation of cytoplasmic α-catenin. This supports a model whereby nesprins could capture cytoplasmic catenins at the nuclear envelope for subsequent nuclear translocation in a force-dependent manner by virtue of a higher affinity in a mechanically relaxed state. Remarkably, this is a distinct mechanism from that proposed for mechanical induction of YAP nuclear translocation whereby nesprins would transmit cytoskeletal forces to stretch nuclear pores open and sterically facilitate nuclear translocation (Elosegui-Artola et al. 2017). The reason may lie in that β-catenin is its own nuclear transporter that directly interacts with nuclear pore proteins (Fagotto, Glück, and Gumbiner 1998). Additionally, our results support that nuclear localization of α-catenin promotes β-catenin nuclear localization by affinity (Fig. 4). However, we show that nuclear α-catenin limits β-catenin-dependent transcription, consistently with previous reports (Giannini, Vivanco, and Kypta 2000; Merdek, Nguyen, and Toksoz 2004; Daugherty et al. 2014; Choi et al. 2013). Since α-catenin needs β-catenin for nuclear localization (Daugherty et al. 2014), we thus propose that β-catenin piggybacks its own retention, and transcription-limiting factor in a nesprin2G tension-dependent manner.

Based on these results and our previous study (Gayrard et al. 2018), we propose that in a manner dependent on the EMT program, mechanosensitive nesprins may capture, at the nuclear envelope, the catenins released from the plasma membrane when E-cadherin relaxes, and thereby fine-tune their nuclear translocation and activities (Fig. 4G).

## Materials & Methods

### Cell lines and culture

Madin–Darby canine kidney (MDCK) type II G and NIH 3T3 cells were cultured at 37 °C and 5% CO_2_ in DMEM supplemented with 10% (vol/vol) FBS with low glucose 1g/L, and 100 µg/mL Hygromycin for stably expressing tagged proteins of interest. Plasmids were transfected using Turbofect according to the instructions of the manufacturer (Thermo Fisher Scientific). Stable cell lines were obtained by FACS after two weeks of selection at 200µg/mL hygromycin (InVivoGen, France). Cell lines expressing β-catenin-GFP and α-catenin-GFP are gifts from W.J. Nelson (Stanford University, Stanford, CA). The cell line expressing TOP-dGFP is a gift from C. Gottardi (Northwestern University, Chicago, IL).

Unless indicated otherwise, cells were plated on glass coverslips coated with 50µg/mL human type IV collagen (Sigma C7521) 24 to 48h before imaging. Before stimulation with HGF, cells were starved 12h in DMEM supplemented with 0.5% FBS. Live cells were imaged in FluoroBrite DMEM medium (Life technology) without phenol red supplemented with 10% or 0.5% FBS depending on experiments, 1U/mL of penicillin, 20 mM Hepes and 2.5mM L-glutamine, at 37 °C and 5% CO_2_.

### Plasmids

TSMod was obtained from the EcadTSMod construct (Borghi et al. 2012) by digestion with BspEI and SpEI.

To generate the CB construct, the cytoplasmic ‘C’ and transmembrane/KASH ‘K’ domains of mN2G were obtained from the mN2G-GFP construct (Luxton et al. 2010) by PCR (fwd(C): CTGGACTAGTGGATCCGAATTCGAGATGGCTAGCCCTGTGCTGCCC, rv(C): CTTTCGAGACTCCGGAGCCTGCTCCTGCTCCTCCACCGGTGTGGGGCATCCTGCTGTCT, fwd(K): GCTGTACAAGACTAGTGGTGCTGGAGGTGGTGCTGTTAACCTCGACAGCCCCGGCAGC, rv(K): TACCGAGCTCGGATCCCTAGGTGGGAGGTGGCCCGT). Digestion and PCR products were cloned into a pcDNA3.1 hygro(-) vector (ThermoFisher Scientific) digested by BamHI using In-fusion HD cloning kit (Clontech Laboratories, TaKaRa Bio Inc., Shiga, Japan).

The CH mutant was made from the CB construct by site-directed mutagenesis (Quikchange II XL, Agilent) (fwd: CCATTATCCTTGGCCTGGCTTGGACCGCTATCCTGCACTTTCATATTG, rv: CAATATGAAAGTGCAGGATAGCGGTCCAAGCCAGGCCAAGGATAATGG).

To generate the SUN2-TSMod construct, the transmembrane ‘T’ and nucleoplasmic ‘N’ domains were obtained from a SUN2-V5 construct (Hodzic et al. 2004) by PCR (fwd(N): CTGGACTAGTGGATCCAAGCTTACCATGGGTAAGCCTATCC, rv(N): CTTTCGAGACTCCGGAGCCTGCTCCTGCTCCTCCACCGGTGTAGCCTGCAAGGTCATCCTCTGA, fwd(T): GCTGTACAAGACTAGTGGTGCTGGAGGTGGTGCTGTTAACACGGACTCAGACCAGCACAG, rv(T): TACCGAGCTCGGATCCCTAGTGGGCAGGCTCTCCGT). Digestion and PCR products were cloned into a pcDNA3.1 hygro(-) vector as above.

The mCherry-DNKASH construct was made from the DNKASH-TSMod construct digested by AgeI/HpaI and mCherry from a PCR on a mCherry-cSrc construct (from M. Davidson, Florida State University, Tallahassee, FL; 55002; Addgene) (fwd: ATTCGAGATGACCGGTATGGTGAGCAAGGGCGAGGAGGATA, rv: GGGGCTGTCGAGGTTAACAGCACCACCTCCAGCACCCTTGTACAGCTCGTCCATGCCG).

The NLS-iRFP-a-catenin construct was made from an alpha-catenin-GFP construct (a gift from W. J. Nelson) digested by AgeI/SalI and iRFP (Filonov et al. 2011) by PCR with the NLS sequence in the PCR primers (fwd: CGCTAGCGCTACCGGTCGCCACCATGCCTGCTGCTAAAAGAGTTAAATTAGATATGGCTGAAGGATCCGTCGCCA, rv: TCATGGTGGCGTCGACTGCAGAATTCGAAGCTTGAGCTCGAGATCTGAGTCCGGACTCTTCCATCACGCCGATCTGC).

Constructs were verified by digestion and gel electrophoresis, and sequencing of coding regions.

The shRNA against a-catenin and lyn-mCherry plasmids were gifts from I.G. Macara (University of Virginia, Charlottesville, VA) and W.J. Nelson, respectively. The mTFP-5aa-Venus and mTFP-TRAF-Venus FRET constructs were gifts from R.N. Day (Indiana University, Bloomington, IN).

### Transient genetic perturbations

Transient expression of NLS-iRFP-a-catenin and mCherry-DNKASH was obtained by transient transfection of the plasmid above using Turbofect according to the instructions of the manufacturer (Thermo Fisher Scientific). Cells were used for experiments between 48 and 72 h after transfection.

Transient Depletion of a-catenin was similarly obtained with a shRNA plasmid co-transfected with a lyn-mCherry plasmid, as a marker for transfected cells. In these conditions, more than 90% of co-transfected cells had a decrease of ∼90% in α-catenin content, as shown previously (Borghi et al. 2010, 2012).

### Chemical inhibitors and biochemical perturbations

Hepathocyte Growth factor (HGF) was used at 50ng/mL final concentration on starved cells (Sigma H5691, 20µg/mL in PBS stock). Cytochalasin D was used at 0.5μM final concentration (Sigma, 10 mg/mL in DMSO stock). Colchicine was used at a 1 µM final concentration (Sigma, 50 mg/mL in Ethanol stock). Y27632 was used at a 10 µM final concentration (Sigma, 10 mg/mL in water stock). EDTA was used at 1.65 µM final concentration (Invitrogen, 0.5M in water stock). LiCl was used at 30 mM final concentration (105679; Merck Chemicals).

### Nuclear confinement

Cells were seeded in a previously described microfluidic device, which exhibited 2,3, and 5 µm-wide, 5µm high constrictions between adjacent 15-30 µm-wide circular Poly-Di-Methyl-Siloxane (PDMS, Sylgard 184, Dow Corning) pillars (Davidson et al. 2015).

### Stretching

As previously described (Fink et al. 2011), collagen lines (10µm wide) were micropatterned on thin silicon membranes (Gel pak PF-60-X4; thickness: 150 μm; Teltek (Sonora, CA)). Membranes were clamped in a custom-made device allowing membrane stretching using a micrometric screw, with a maximal extension of ∼ 25% in ∼ 30 s. A rectangular polydimethylsiloxane 300 µL chamber was attached onto the membrane using vacuum grease, and cells were seeded for 24 h before stretching. FRET measurements were within seconds before stretch and approximately 1min after. Cell and nucleus strains are percent increases in length along the stretch axis.

### Wound Healing and confined collective migration

Wound healing: cells were cultured at confluence around a 5*5mm PDMS stencil. The PDMS stencil was removed 24h after cell seeding and 48 before imaging.

Confined collective migration: cells were cultured at confluence around a PDMS slab exhibiting 100 µm-high, 40 µm-wide micromolded channels in contact with the coverslip. Cells were imaged 24 to 48h after seeding.

Cell velocities were averages of instantaneous velocities (between consecutive frames) of each cells.

### FRET Imaging

Spectral imaging was performed on a confocal microscope (Carl Zeiss LSM 780) with a 63x/1.4NA Plan-Apochromat oil immersion objective. mTFP1 was excited by the 458 nm line of a 25-mW argon laser. Emission was sampled at a spectral resolution of 8.7nm within the 476– 557nm range on a GaAsP detector. For time lapse experiments (nuclear confinement, wound healing and confined collective migration) images were acquired every 10min during ∼15 hours for ∼10 different positions.

### FRET analysis

Fluorescent images were analyzed in Image J using the Fiji distribution and the publicly available PixFRET plugin. All channels were background-subtracted, Gaussian smoothed (radius=1pixel) and thresholded (above ∼3-5% of the 12bits range). The FRET index *E*_R_ was computed pixel-by-pixel as *I*_EYFP_/(*I*_mTFP_+*I*_EYFP_), where *I*_mTFP_ and *I*_EYFP_ are the intensities in 494nm and 521nm channels. Unless specified otherwise, the FRET index was then averaged over the segmented nuclear envelope for comparison between conditions. The FRET index did not display a significant dependence on the z section considered (Fig. S3B). FRET efficiencies *E* were computed from FRET indices *E*_R_ with *E*=(1-*a*(1-*E*_R_))/(1-*b*(1-*E*_R_)), where *a* and *b* account for donor spectral bleed-through, acceptor direct excitation, and differences in donor vs acceptor absorption cross-sections and detection efficiencies (Lee et al. 2005). Measured FRET indices *E*_R,H_ and *E*_R,L_ (Fig S3A), and published FRET efficiencies *E*_H_ and *E*_L_ of mTFP-5aa–Venus and mTFP-TRAF-Venus constructs (Day, Booker, and Periasamy 2008) were used to recover *a*=(*E*_H_(1-*E*_R,H_) - *E*_L_(1-*E*_R,L_) + *E*_H_*E*_L_(*E*_R,H_ - *E*_R,L_))/*c* and *b*= (*E*_H_(1-*E*_R,L_) - *E*_L_(1-*E*_R,H_) + *E*_R,L_ - *E*_R,H_)/*c*, with *c*=(*E*_H_ - *E*_L_)(1 - *E*_R,H_)(1 - *E*_R,L_). Previously published FRET efficiency to force calibration (Grashoff et al. 2010) was then used to retrieve forces, with the CH mutant as the zero-cytoskeletal-force reference.

### Immunostaining

Cells were fixed in 4% PFA (Electron Microscopy Science) for 15 min at RT, permeabilized with 0.5% Triton X100 in PBS for 5 min at 4°C, incubated with 50mM NH4Cl in PBS and blocked for 30 min at RT with 1% BSA (Jackson Immunology) in PBS. Cells were stained with anti-nesprin 2 rabbit antibody directed against CH domain of Nesprin 2 (gift from G.G. Gundersen, Columbia U, New York, NY (Luxton et al. 2010)) (1/500) in PBS 1%BSA for 2h at RT, followed by incubation with Dylight 650 secondary antibody (1/500) (ThermoFisher) for 45 min at RT. Coverslips were mounted in Fluoromount medium (Sigma).

### Fluorescence imaging

Non-FRET fluorescence imaging was performed either on a confocal microscope (Carl Zeiss LSM 780) with a 63x/1.4NA oil immersion objective or on a widefield microscope (Carl Zeiss Axio Observer Z.1) with a 25x/0.8NA oil immersion objective. On the confocal, fluorophores were excited with the 488nm line of the argon laser, the 561nm line of the 15mW DPSS laser and the 633nm line of the 5mW HeNe laser, emmission was collected between 498-561nm, 570-650nm and 638-755nm on PMT detectors. On the widefield, fluorophores were excited with a LED lamp (CoolLED pE-300 white) by using BP excitation filters 450-490nm, 538-562nm and 625-655nm and acquired through BP emissions filters 500-550nm, 570-640nm and 665-715nm on a sCMOS camera (OrcaFlash4 LT, Hamamatsu).

### Fluorescence analysis

Fluorescent images were analyzed in ImageJ using the Fiji distribution. All channels were background-subtracted. Mean pixel intensities were measured within the boundaries of regions of interest (cytoplasm, nucleus and nuclear envelope, which was segmented from the anti-Nesprin 2 antibody or mCherry-DNKASH signals). Nucleus/Cytoplasm ratios were computed from mean intensities. Normalized intensities lie between the minimum (0) and maximum (1) intensities for each condition (HD, LD…).

### Statistical Analysis

Data are presented as mean ± SEM. P-values were calculated from unpaired, non-parametric, two-tailed tests (Mann–Whitney or Krustal-Wallis depending on condition numbers) in GraphPad Prism V software. n.s.: statistically non-significant. ****:<0.0001, ***:<0.001, **:<0.01, *<0.05. R were calculated from linear regression by the least-square method.

## Acknowledgements

We thank V. Doye, M. Piel and the members of the laboratory for insightful discussions. We thank W.J. Nelson (Stanford University, Stanford, CA), C. Gottardi (Northwestern University, Chicago, IL). I.G. Macara (University of Virginia, Charlottesville, VA), R.N. Day (Indiana University, Bloomington, IN) and G.G. Gundersen (Columbia University, New York, NY) for sharing reagents.

This material is based upon work supported by the Centre national de la recherche scientifique (CNRS), the Agence nationale de la recherche (ANR) grants ANR-13-JSV5-0007 and ANR-14-CE09-0006, France BioImaging (ANR-10-INBS-04), La Ligue contre le Cancer (REMX17751 to PMD) and the Fondation ARC (PDF20161205227 to PMD). PSC has received funding from the European Union’s Horizon 2020 research and innovation programme (Marie Skłodowska-Curie grant agreement #665850-INSPIRE) and acknowledges the Ecole Doctorole FIRE-Programme Bettencourt. ERG was supported by a European Research Council consolidator grant.

We acknowledge the ImagoSeine core facility of the Institut Jacques Monod, member of IBiSA and France-BioImaging (ANR-10-INBS-04) infrastructures.

## Author contributions

TD and PSC performed experiments together with PMD (Nuclear confinement) and DC (Stretching). TD, PSC, PMD, CS, DC, BC, ERG, and NB contributed reagents. TD, PSC and NB analyzed the data. NB wrote the manuscript. All authors discussed the results and commented on the manuscript.

## Conflif of interest

The authors declare no conflict of interest.

---

**Figure S1.**
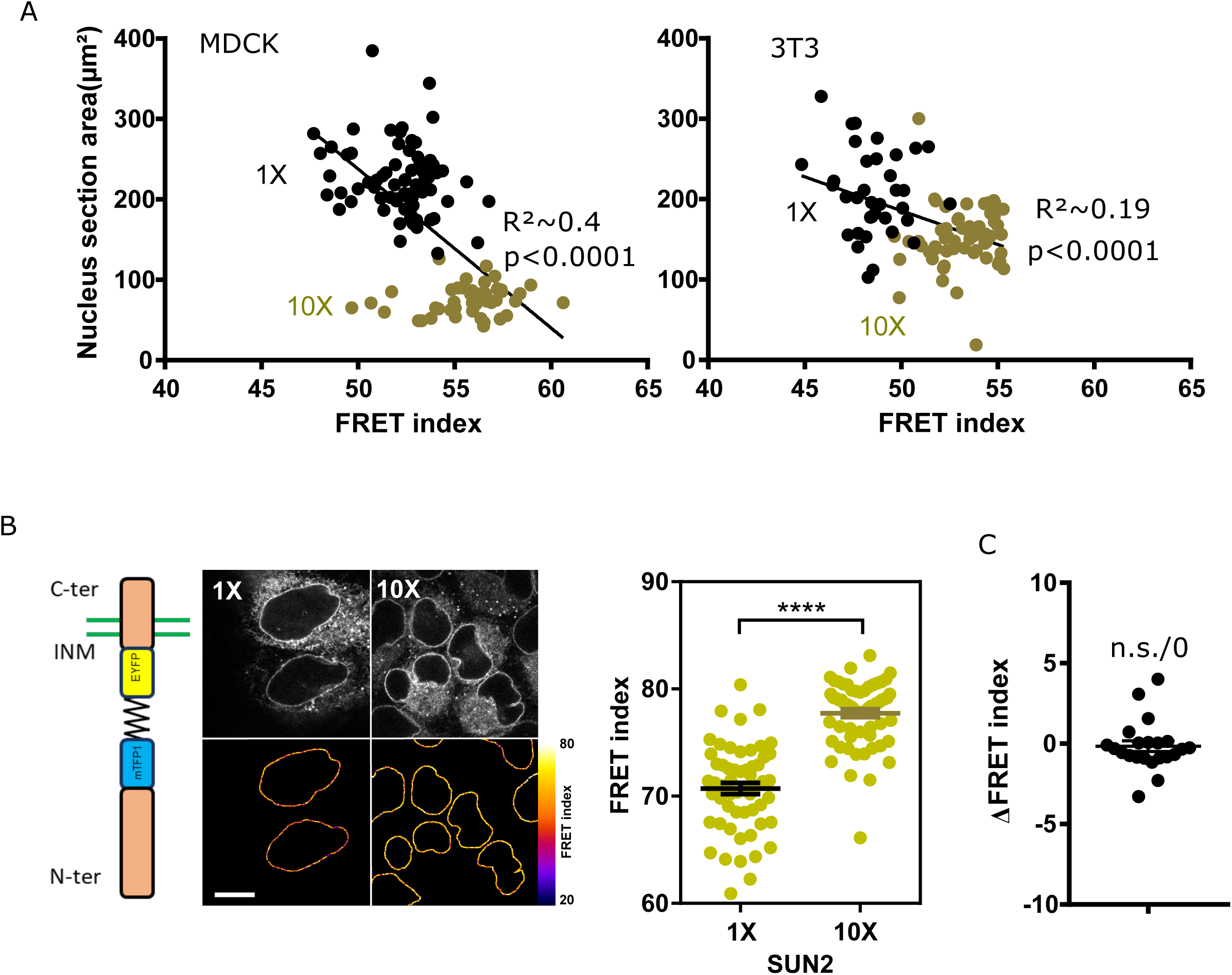
A) Nucleus section area as a function of FRET index for low (1X) and high (10X) cell densities for MDCK (left, n= 73 1X, 45 10x) and NIH 3T3 (n=36 1X, 59 10X) cells. Solid lines are linear fits. B) Left: sketch of the SUN2-TSMod construct. Middle: MDCK cells expressing the SUN2-TSMod construct plated at 5.10^2^ cells/mm^2^ (1X) and 5.10^3^cells/mm^2^ (10X). Right: FRET index of the SUN2-TSMod construct at 1X and 10X densities, in MDCK cells (n= 58 1X, 62 10X). C) FRET index difference between the front and back of a nucleus within leader cells (n= 22). Bar=5 µm. Mean ± SEM. Kruskal-Wallis test.

**Figure S2.**
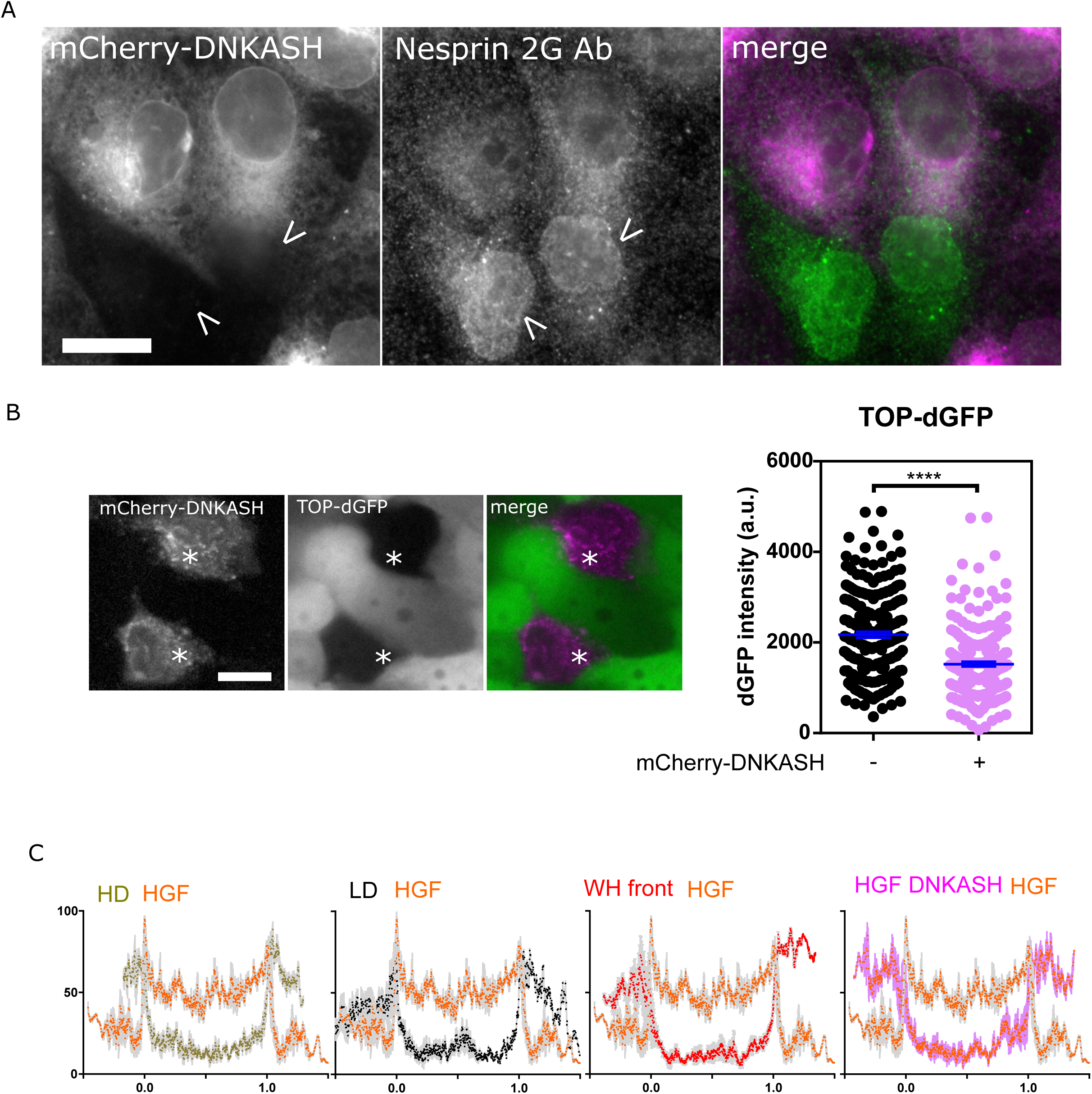
A) MDCK cells transiently expressing mCherry DNKASH and stained for Nesprin2G. Only non-transfected cells (arrow head) show Nesprin2G localization at the nuclear envelope. B) MDCK cells stably expressing TOP-dGFP and transiently expressing mCherry-DNKASH after 10hrs with LiCl. Cells expressing mCherry-DNKASH (star) show lower dGFP levels (n= 216 +mCherry-DNKASH, 216 - mCherry-DNKASH). C) Normalized a-catenin-GFP intensity along a linescan across the nucleus of cells exposed to HGF compared to that of cells plated at high (HD, 10X) and low (LD, 1X) densities, at the front of an epithelial wound, and expressing mCherry-DNKASH with HGF. Linescans are averages of 3 cells, 5 pixels-window moving-average. 0 and 1 are nuclear envelope positions. Bar=10 µm. Mean ± SEM. Kruskal-Wallis test.

**Figure S3.**
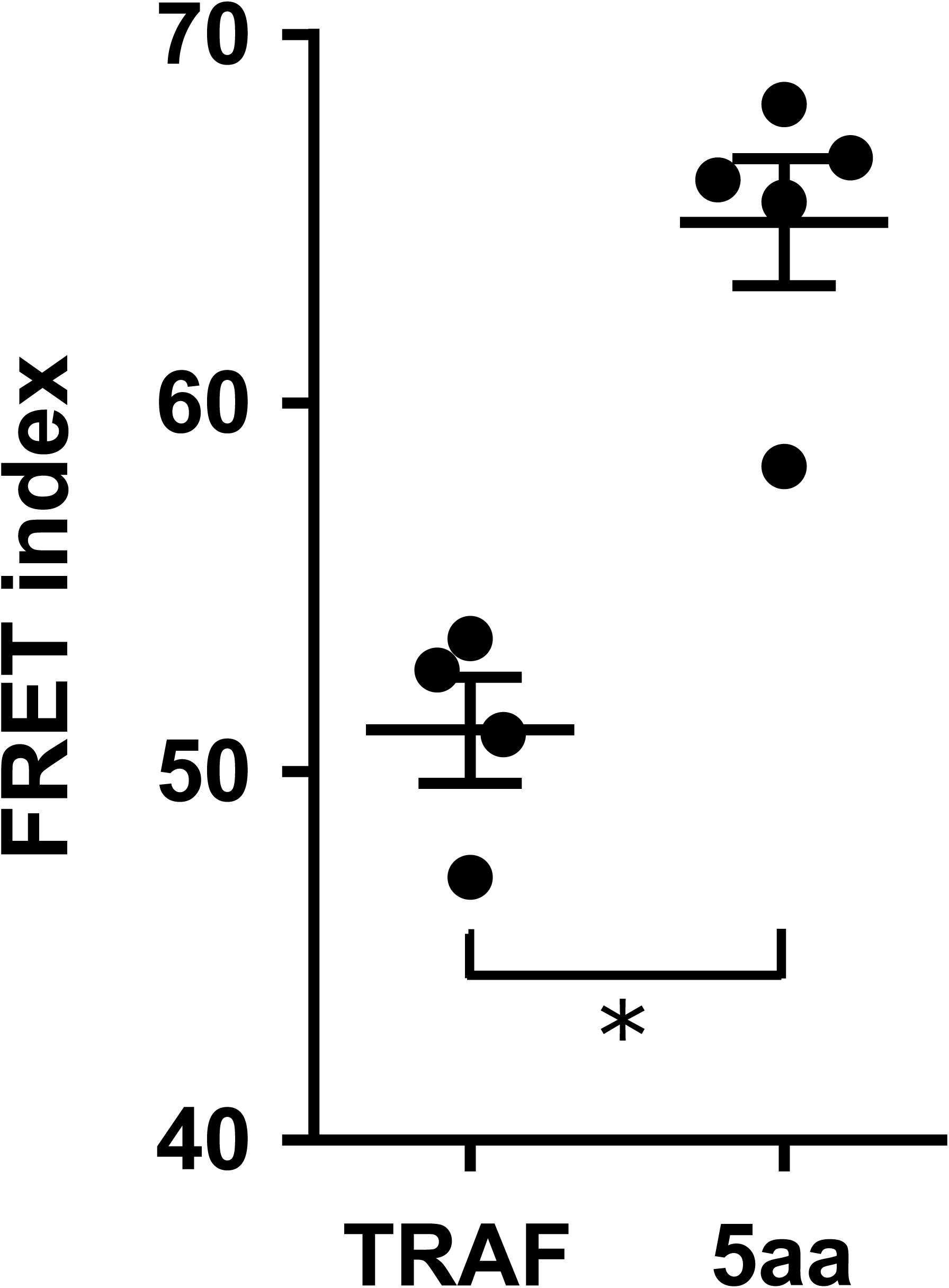
A) FRET index of 5aa and TRAF standards expressed in MDCK cells (n=4). B) Left: sketch of a cell growing on the side of a PDMS channel for FRET analysis along z. Direct fluorescence and FRET index of an MDCK cell stably expressing the CB construct from the region in the dotted box above. Right: normalized FRET index from the white dotted boxes on the left (n=4). Mean ± SEM. Kruskal-Wallis test.

